# The microbiome of Total Suspended Particles (TSP) and its influence on the respiratory microbiome of healthy office workers

**DOI:** 10.1101/2024.08.12.607611

**Authors:** Giulia Solazzo, Sabrina Rovelli, Simona Iodice, Matthew Chung, Michael Frimpong, Valentina Bollati, Luca Ferrari, Elodie Ghedin

## Abstract

Air particulate matter (PM) is widely recognized for its potential to negatively affect human health, including changes in the upper respiratory microbiome. However, research on PM-associated microbiota remains limited and mostly focused on PM (e.g., PM_2.5_ and PM_10_). This study aims to characterize for the first time the microbiome of Total Suspended Particles (TSP) and investigate the correlations of indoor TSP with the human upper respiratory microbiome. Biological and environmental samples were collected over three collection periods lasting three weeks each, between May and July 2022 at the University of Milan and the University of Insubria Como. TSP were sampled using a filter-based technique, while respiratory samples from both anterior nares (AN) and the nasopharynx (NP) were collected using swabs. Microbiome analysis of both human (N = 145) and TSP (N = 51) samples was conducted on metagenomic sequencing data. A comparison of indoor and outdoor TSP microbiomes revealed differences in microbial diversity and taxonomic composition. The indoor samples had higher relative abundance of environmental bacteria often associated with opportunistic infections like *Paracoccus* sp., as well as respiratory bacteria such as *Staphylococcus aureus* and *Klebsiella pneumoniae*. Additionally, both indoor and outdoor TSP samples contained broad spectrum antibiotic resistance genes. Indoor TSP exposure was negatively associated with commensal bacteria and positively associated with *Staphylococcus aureus* relative abundance. Finally, a correlation between the relative abundance of respiratory bacteria identified in the indoor TSP and the upper respiratory microbiome was found, suggesting a potential interaction between TSP and the upper airways.

## 1. Introduction

Particulate matter (PM) represents a group of air pollutants consisting of inhalable solid and liquid particles suspended in the atmosphere. PM is classified based on the aerodynamic diameter of particles, with ultrafine particles having the smallest diameter (less than 0.10 μm, PM_0.1_), and Total Suspended Particles (TSP) being the largest, with an aerodynamic diameter of less than 100 μm. These particles are common in both outdoor and indoor environments, and could lead to unhealthy exposures if concentrations exceed established thresholds (Bai et al., 2018). Numerous studies have demonstrated the association between both short- and long-term exposures to these pollutants with respiratory and cardiovascular diseases (Bezirtzoglou et al., 2011; Palacio et al., 2023; Peden, 2024). The biological mechanisms behind this association are not fully understood, but many studies have observed that exposure to PM induces oxidative stress, inflammation, epithelium damages and immune dysregulation in the upper airways (Xue et al., 2020). Despite the well-known adverse effects of PM exposure on the human respiratory system, few studies have characterized its microbiota, and with a primary focus on microbial allergens and pathogens in PM_2.5_ and PM_10_ (Madsen et al., 2023; Qin et al., 2020; Zhou et al., 2021). Moreover, these studies have centered on either the outdoor or the indoor PM microbiota but have not assessed the differences in the microbial communities between these two types of exposures. In addition, because people usually spend 80-90% of their time in indoor spaces (such as homes, schools, offices, and gyms, for example) a better characterization of these environments could help inform prevention policies (Bennitt et al., 2021; Li et al., 2023; Peixoto et al., 2023; Wang et al., 2023).

Given that adults typically inhale, on average, 6-7 liters of air per minute, determining what microbes the human upper respiratory airways are exposed to via suspended particulate matter is important (Gisler et al., 2021; Hamidou Soumana and Carlsten, 2021; Li et al., 2019; Mariani et al., 2021; Rylance et al., 2016; Xue et al., 2020). The upper respiratory microbiome, as one of the first layers of interaction with inhalable pollutants, plays a crucial role in maintaining host health (Hou et al., 2022). Although the number of studies examining the effects of PM exposure on the respiratory microbiome is limited, data suggest that this exposure can impact commensal bacteria and microbial diversity (Vieceli et al., 2023). For example, two studies have shown that exposure to PM_10_ and PM_2.5_ is associated with a decrease in commensal bacteria present as part of the nasal microbiota (Mariani et al., 2018; Padhye et al., 2021), and an increase in the abundance of potential pathogens such as *Moraxella*, *Haemophilus*, and *Staphylococcus* (Mariani et al., 2018; Qin et al., 2019).

In this study, we characterized for the first time the indoor TSP microbiome collected in an office setting across 2 campuses in Italy and compared it to the corresponding outdoor TSP microbiome to investigate the effects of the outdoor environment on indoor microbial diversity. We also assessed whether correlations could be determined between the indoor TSP and microbial communities of the upper respiratory tract of healthy individuals by characterizing the microbiome of their anterior nares and nasopharynx.

## 2. Methods

### 2.1 Recruitment and Samples collection

Healthy volunteers employed by the University of Milan and the University of Insubria, Como, were enrolled in the study and sampled within their workplace over 3 weeks between May and July 2022. Each week, we sampled both indoor and outdoor TSP. The indoor TSP sampling was carried out in 13 different offices located in 4 buildings (two at the University of Milan and two at the University of Insubria, Como). Eligibility criteria for subject enrollment included: age between 20-65 years old with no known chronic or infectious disease, working ≥ 10 hours per week, and ≥ 2 days per week spent in the office. Each participant was asked to fill out a questionnaire of relevant information over the course of each week, such as mask-wearing in the office, number of cigarettes smoked, transport used for commuting to work, sports and other activities performed each week. We enrolled 26 subjects and assigned each of them to a monitoring group (MG) defined by when the TSP was monitored. The first group includes subjects (N = 8) from the Milan site monitored from the 9^th^ to the 27^th^ of May (monitoring group 1 = MG-1); the second group includes other subjects (N = 8) from Milan monitored from the 6^th^ to the 24^th^ of June (monitoring group 2 = MG-2); finally, the third group includes subjects (N = 10) from the Como site monitored from the 11^th^ to the 29^th^ of July (monitoring group 3 = MG-3). Subject characteristics are described in **Table 1**. Most of the participants were non-smokers who live in a city or a small town with moderate traffic (**Table 1**).

**Table 1.** Healthy volunteers characteristics.

| **Table 1. Healthy volunteers characteristics** |  |
| --- | --- |
| **Characteristics** | **(N= 26)** |
| ***Sex, n (%)*** |  |
| Female | 17 (65.4%) |
| Male | 9 (34.6%) |
| ***Age, mean (SD)*** | 38.8 (8.7) |
| ***BMI, mean (SD)*** | 24.3 (3.2) |
| ***Smoking, n (%)*** |  |
| No | 19 (73.1%) |
| Yes | 7 (26.9%) |
| ***Days in office per week, mean (SD)*** | 4.4 (0.8) |
| ***Wearing face mask in the office, n (%)*** |  |
| No | 21 (81%) |
| Yes | 5 (19%) |
| ***Residential area, n (%)*** |  |
| City | 12 (46.2%) |
| Small town | 13 (50%) |
| Rural area | 1 (3.8%) |
| ***Traffic level in residential area, n (%)*** |  |
| High | 7 (27%) |
| Medium | 16 (62%) |
| Low | 3 (12%) |

At the end of every working week, anterior nares and nasopharynx swabs were collected. By the end of the study, a total of 145 samples were collected from the 26 individuals enrolled. Concurrently, indoor and outdoor TSP samples were collected weekly using an active filter-based technique comprised of a low-flow air SKC sampling pump attached to a filter cassette (SKC inc.) containing a 37-mm, 0.8 µm mixed cellulose esters (MCE) membrane filter (Omega, SKC Inc.). In each office, the active sampling line was placed at the center of the room, at a minimum distance of 1 m from the wall and away from ventilation channels and heating sources, with the filter cassette at a height corresponding approximately to the breathing zone of seated occupants. Furthermore, TSP were simultaneously sampled at two outdoor sites, that remained constant, chosen to be as close as possible to the offices to obtain the best possible estimates of TSP outdoor concentrations over the 3 monitoring groups (MG-1, MG-2, MG-3). For both indoor and outdoor monitoring, the sampling pumps were set at a flow ranging from 2 to 4 L/min and each sampling lasted 8 hours per day, for 5 consecutive days. This study was approved by the Ethics Committee of the University of Milan (approval number 24/22), in agreement with the principles of the Helsinki Declaration. All participants signed a written informed consent.

### 2.2 Total Suspended Particles (TSP) mass measurements

The TSP mass of each filter was determined gravimetrically, following a weighing standard operating procedure. Briefly, before weighing, each filter was conditioned for 48h at a constant temperature of 20°C ± 1°C and 50% ± 5% relative humidity values in a controlled environment (SCC 400L Climatic Cabinet, Sartorius, Varedo (MB), Italy). After this step, the filter was weighed three times using a micro-balance (readability of 1 μg) (GIBERTINI 1000, Novate, Milan, Italy), which was regularly checked for calibration using certified standard weights of 100 and 1000 mg. An electrical C-shaped ionizer (HAUG GmbH & CO. KG, Germany) was used to eliminate electrostatic charges from the filter surface. This procedure was repeated before and after each sampling and the TSP mass was determined by differential weighing. Two laboratory blanks were weighed along with the filters under the same conditions to verify possible anomalies in the weighing room conditioning and the average blank filter masses were then used to correct the filter mass results for each test. After the mass measurements, the collected filters were stored at −80°C until further analysis.

Since negligible differences were found between the TSP outdoor concentrations measured at the two outdoor sampling sites, both in Milan and Como, the outdoor exposure was estimated using the mean of the two measurements performed in each building every week. A Wilcoxon signed-rank test was used to analyze possible differences between indoor and outdoor TSP concentrations. These analyses were performed using R software v 4.2.1.

### 2.3 Whole genome shotgun (WGS) sequencing and bioinformatics analyses

The DNA from the cellulose filters was extracted using the NucleoSpin cfDNA XS kit (Macherey-Nagel), while the DNA from swabs (anterior nares and nasopharynx) was extracted using the QIAamp UCP Pathogen Mini (Qiagen, Hilden, Germany) protocol. All DNA samples were quantified using the Qubit dsDNA High Sensitivity Assay kit (Invitrogen). The sequencing paired-end library was prepared using the NexteraXT DNA library preparation kit (Illumina) following the manufacturer’s protocol. To purify the library, the Agencourt AMPure XP beads (Beckman Coulter, Inc.) was used. Finally, all barcoded samples were combined to create one equimolar pool for sequencing. At this point, the sequencing pool was quantified and sequenced on the Illumina Novaseq 6000 S2 in one single run (sequence length 150 bp, paired end). The reads obtained from the whole genome shotgun (WGS) sequencing were demultiplexed using Bcl2fastq (v 2.20) and trimmed using fastp (v 0.20.1). Taxonomy was assigned using Kraken (v 2.1.2) and Bracken (v 2.5) (Lu et al., 2017; Wood and Salzberg, 2014). Host reads were filtered from the FASTQs by removing any reads mapping to a reference human genome (GrCh38) using Bowtie (v 2.4.2) (Langmead et al., 2009). The processed reads were given taxonomic assignments using a database consisting of human, bacterial, fungal, archaeal, and viral RefSeq databases along with the EuPathDB46 database. The antimicrobial resistance profile was analyzed using the MEGARes database (v 3.0) (Lakin et al., 2017). The *Decontam* R package (v 1.18) was used to identify contaminants from all samples in the dataset using as reference three types of negative controls: sampling (clean swabs opened at the sampling site); DNA extraction (all the reagents plus water); library preparation (all reagents plus water). The alpha (α) diversity was calculated using the Shannon Index and the beta (ß) diversity using the Bray-Curtis matrix. To calculate and analyze both the alpha and beta diversities we used the following R packages: *phyloseq* (v 1.42), *microbiome* (v 1.22), *microViz* (v 0.10), and *MicrobiotaProcess* (v 1.10) (McMurdie and Holmes, 2013; Xu et al., 2023). The non-parametric paired t-test was used to analyze the difference in alpha diversity among groups. A Spearman correlation test was performed to check the correlations of the indoor TSP concentrations with the alpha diversity of the indoor TSP microbiome and the respiratory microbiome. The difference in beta diversity was analyzed with the Analysis of Similarities (ANOSIM) and the permutational multivariate analysis of variance (PERMANOVA) with 999 permutations using adonis2 (Anderson, 2001) from *MicrobiotaProcess* (v 1.10). The ANOVA-like Differential Expression tool (ALDEx2) and the Analysis of Compositions of Microbiomes with Bias Correction (ANCOM-BC) for differential abundance analysis (Lin and Peddada, 2020) were then used to test for the difference between environmental samples (indoor vs outdoor TSP) and between respiratory samples (anterior nares vs nasopharynx). We considered an association significant when a p-value < 0.05 and FDR < 0.10 were reached. The composition of the microbiome was described using the *microViz* R package (v 0.10). The association between indoor TSP exposure and the microbiome was analyzed by multivariable association using the *nlme* package (v. 3.1). Bacterial abundance was determined using the centered log-ratio (CLR) transformation. The model was adjusted for age, gender, days spent in the office, and habitual use of a face mask. The sample observations were grouped by “*subject id”*. Finally, we selected the skin/respiratory bacteria identified among the top 200 bacteria found in the indoor TSP microbiome and we determined the correlation of their relative bacteria abundance between the indoor TSP and the respiratory samples. The correlation and the plots were generated using the R package *ggstatsplot* (v. 0.12.2). We also tested for relative abundance correlations among the top antimicrobial resistance classes comparing observations in the indoor TSP samples with the respiratory microbiome. All statistical analyses were performed with R software, v 4.2.1.

## 3. Results

### 3.1 Microbiome differences between indoor and outdoor TSP

To quantify TSP concentrations and their potential association with microbial composition of the ambient air inside offices and outside the buildings where these offices are located, we collected 37 indoor and 17 outdoor TSP samples on 2 university campuses over 3 sampling periods. We first measured TSP concentrations and analyzed the differences between the cities (Milan and Como), between the types of exposures (indoor and outdoor), and across the monitoring groups (MG-1, MG-2, MG-3). We observed that the offices located in Milan had higher indoor TSP concentrations (Wilcoxon, p-value < 0.001) compared to the offices located in Como. Specifically, the first group in Milan (MG-1) had the highest indoor TSP compared to the second group in Milan (MG-2) and the one in Como (MG-3) (**Table 2)**. Significant differences in indoor TSP concentrations were observed across all three monitoring groups (**Supplemental Figure 1)**. As for the indoor exposure, the outdoor TSP concentrations were significantly higher in Milan than in Como (Wilcoxon, p-value = 0.04). Among the three monitoring groups, the difference was significant (Wilcoxon, p-value = 0.04) only for outdoor concentrations compared between MG-1 and MG-3. Finally, the indoor TSP concentrations were significantly lower (Wilcoxon, p-value = 0.002) than the outdoor TSP concentrations (**Table 2)**. This general trend was observed across the sampling weeks with a calculated ratio between the indoor and the corresponding outdoor TSP concentrations (I/O ratios) generally below 1 **(Supplemental Table 1)**.

**Table 2.**
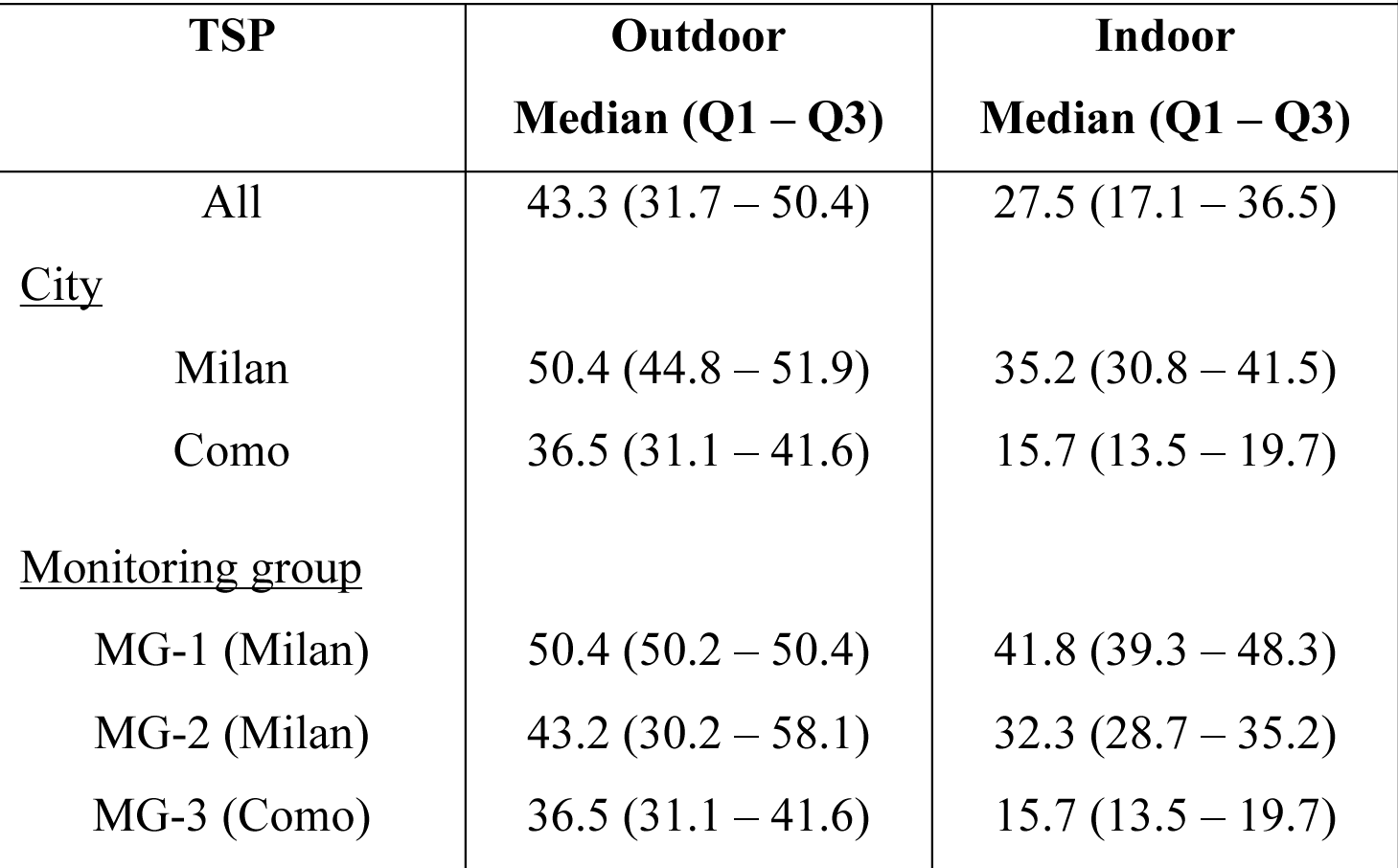
Description of indoor and outdoor TSP levels (µg/m^3^)

Since the indoor TSP differed significantly from the outdoor TSPs, we tested whether this also applied to the composition of the microbiome present on the particles. For this, we performed a metagenomic analysis by whole genome shotgun sequencing. We excluded from the analysis samples with sequencing depths below 100,000 reads. We obtained 51 TSP samples with a median sampling depth of 385,900 sequence reads per sample (range: 103,408 – 2,198,650 reads). After taxonomic assignments of the sequenced reads, we compared the microbial diversity of the indoor samples collected across 13 offices and the correlation with TSP concentrations. While we did not find significant differences in beta diversity (which compares microbial community topology) among the monitoring groups, we did observe a positive correlation between alpha diversity—a measure of taxonomic richness and relative abundance of a sample—and TSP concentrations (Spearman correlation test, rho = 0.5, p-value = 0.001). In addition, we found that the monitoring group that had the highest indoor TSP concentrations (MG-1) had an alpha diversity significantly different from the other two monitoring periods (**Figure 1)**.

**Figure 1.**
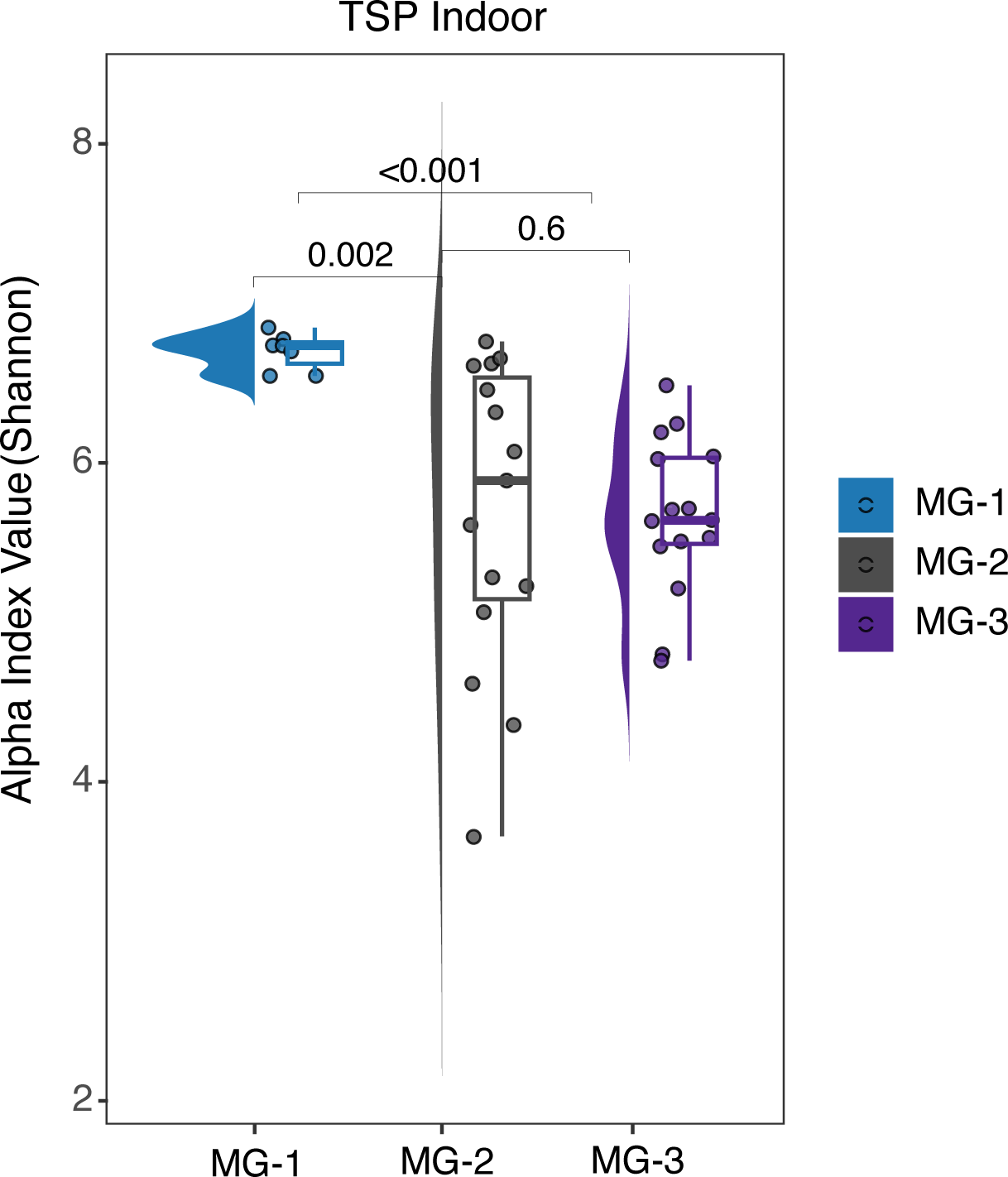
Comparison of alpha diversity of the *indoor* TSP samples across the three monitoring groups (MG-1, MG-2, MG-3). The alpha diversity was estimated using the Shannon Index and the comparison was performed with the Wilcoxon test.

The alpha diversity of the outdoor samples was significantly different between the first monitoring group (MG-1) and the third monitoring group (MG-3) (Wilcoxon test, p-value = 0.01), Specifically, it was higher in MG-1 (Shannon Index, median = 6.7; Q1 – Q3 = 6.6 – 6.7) than MG-3 (Shannon Index, median = 5.1; Q1 – Q3 = 4.8 – 5.7). In line with the outdoor TSP concentrations that were significantly higher in MG-1 compared to MG-3 (**Table 2)**. Finally, there were no significant differences in beta diversity among the three monitoring groups (MG-1, MG-2, and MG-3). When comparing the indoor and outdoor TSP samples, no significant differences in alpha diversity were found across all samples, but a significant difference in beta diversity (ANOSIM, R = 0.25, p-value = 0.01) was observed. This difference explained approximately 4% of the diversity (PERMANOVA, p-value = 0.01) (**Figure 2A**). The relative abundance of the top 10 taxa found across the indoor and outdoor TSP samples are described in **Figure 2B**. In both indoor and outdoor TSP samples, the most abundant bacteria were two gram-positive species commonly found in the environment, *Micrococcus luteus* and *Kocuria rhizophila* (**Figure 2B**) and previously described in some air pollutants (Kooken et al., 2012; Madsen et al., 2023). Most of the bacteria found to be differentially abundant were from soil, dust, or plants (e.g., *Xanthomonas campestris* and *Clavibacter michiganensis*), with a few also found to be associated with human skin (e.g., *Micrococcus luteus, Paracoccus yeei*, and *Kocuria rosea*). Differential abundance analyses using two different tools, ALDEx2 and ANCOM-BC, indicate that the relative abundance of several human-associated microbes such as *Paracoccus* species (e.g., *P. contaminans*, *P. yeei*, *P. sanguinis*, *Paracoccus sp. MC1862*), *Moraxella osloensis* and *Corynebacterium diphtheriae* was significantly higher in the indoor TSP compared to the outdoor TSP **(Supplemental Figure 2)**.

**Figure 2.**
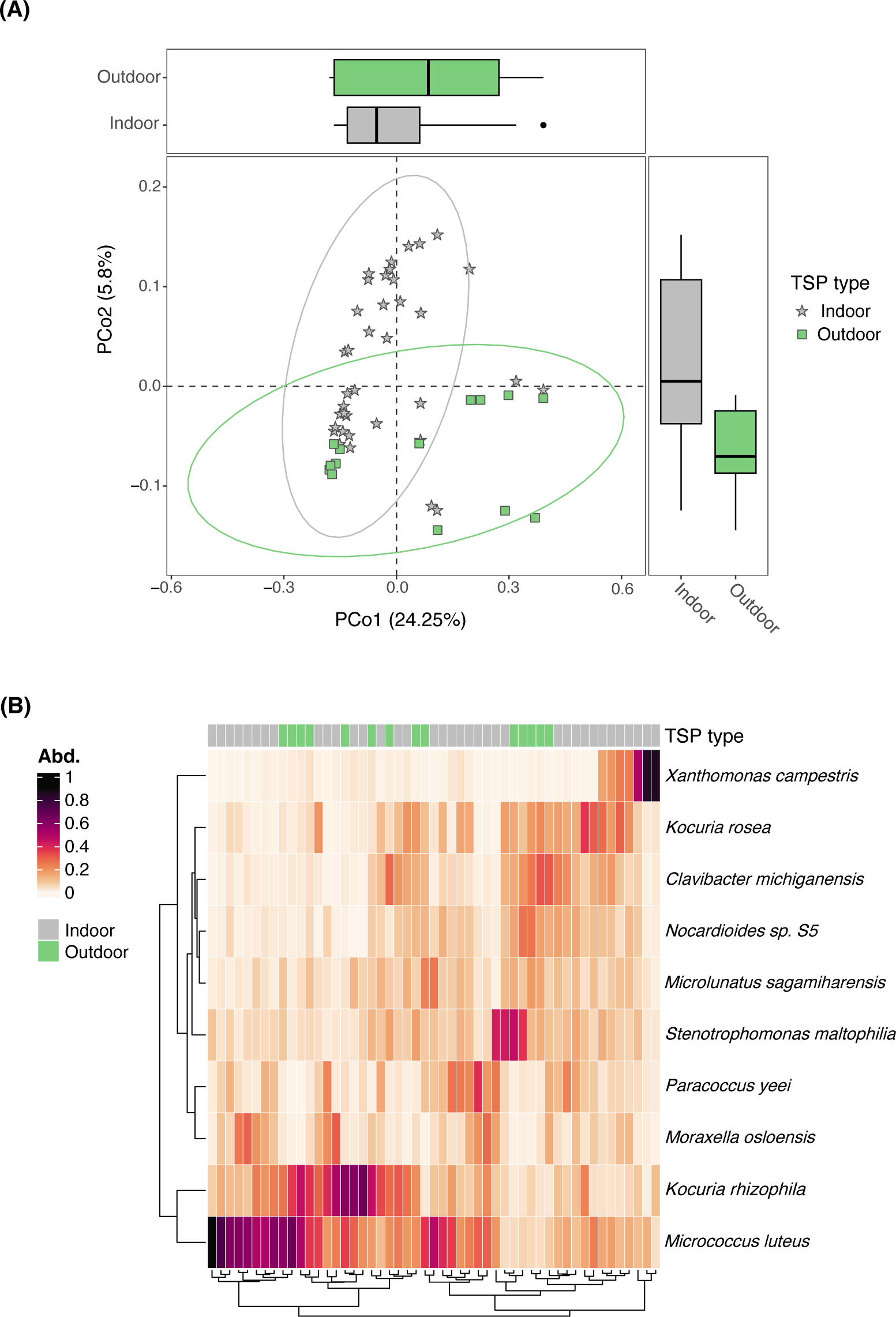
Difference in microbial diversity and composition between indoor and outdoor TSP. (**A)** Difference in beta diversity between indoor (grey stars) and outdoor (green squares) samples. The difference was calculated using the principal coordinate analysis (PCoA) and the Bray-Curtis matrix. The y-axis and the x-axis represent the two main coordinate axes, the percentages are the percent of total variation explained by each axis. (**B**) Heatmap of the top 10 bacterial species across all the TSP samples (indoor and outdoor) and their relative abundance (Abd.).

Metagenomics data also contain functional information, such as antibiotic resistance genes that could be carried by the bacteria. We assigned sequence reads to 57 classes of antibiotics using the antimicrobial resistance database MEGARes (Lakin et al., 2017). While we did not find significant differences in antimicrobial resistance between the indoor and outdoor TSP groups, we did observe high relative abundance of genes associated with resistance to broad-spectrum antibiotics, such as tetracycline and metals (**Figure 3)**.

**Figure 3.**
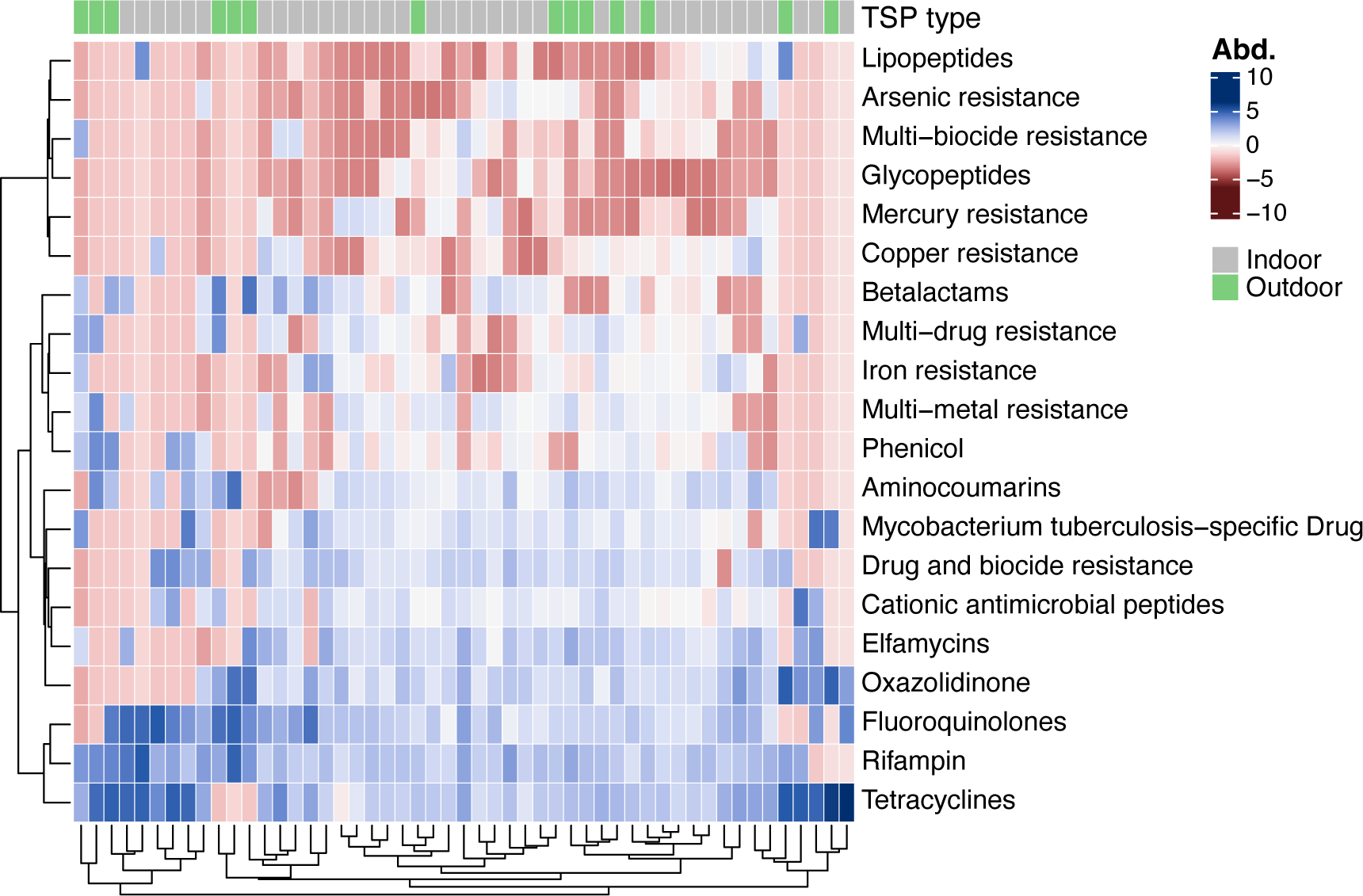
Heatmap of the top 20 antimicrobial chemical classes identified in both outdoor and indoor TSP samples using shotgun sequencing data and the MEGARes (v3.0) database. Each row represents a distinct class, and the color indicates the relative abundance of sequence reads that map to the class of antibiotics for each sample. The abundance was center-log ratio (CLR) transformed.

### 3.2 Correlations in microbial composition between upper respiratory samples and indoor TSP

To assess a possible relationship between the indoor TSP and the upper respiratory microbiome, we characterized the microbial composition of samples collected from the anterior nares (AN) and the nasopharynx (NP) of the subjects, and tested for correlations between microbial composition and TSP concentrations. From the 26 recruited subjects in our study, we collected and sequenced 145 respiratory samples: 73 AN swabs (median sequencing depth = 32,192,366 reads, range: 15,169,918 – 74,149,086), and 72 NP swabs (median sequencing depth = 26,430,743 reads, range: 4,302,478 – 41,851,762 reads). The top 10 bacterial species across all the samples were mostly gram-positive bacteria that are usual commensals of the upper airways (**Supplemental Figure 3**). The most abundant bacteria include several *Corynebacterium* sp. (i.e., *C. propinquum*, *C. segmentosum*, *C. accolens*), *Cutibacterium acnes*, *Dolosigranulum pigrum*, *Staphylococcus aureus* and *Staphylococcus epidermidis*. The alpha diversity was higher in the NP microbiome compared to the AN (Wilcoxon test, p-value < 0.001). The beta diversity was also significantly different between the two types of samples (ANOSIM, R = 0.61, p-value = 0.001) and explained around 13% of the diversity (PERMANOVA, R^2^ = 0.13, p-value < 0.001) **(Supplemental Figure 4)**. While the bacterial composition for the top 10 taxa was similar between the two types of samples, differential abundance analyses indicate significant differences for several bacteria. Among the top 10 bacteria, the AN microbiome had higher relative abundance of skin bacteria such as *Cutibacterium acnes*, *S. epidermidis*, and several *Corynebacterium* (e.g., *C. kefirresidentii*, *C. segmentosum*, *C. accolens*, *C. macginleyi*), while the NP had higher relative abundance of *Staphylococcus aureus* **(Supplemental Figure 5)**.

We analyzed the association between microbial taxa present in the microbiome from each sample type with indoor TSP exposure. For the AN, the indoor TSP exposure was negatively associated with the relative abundance of *Streptococcus pneumoniae* (beta = −0.015, 95% CI = 0.7 – 0.9, p-value = 0.01), but positively associated with the relative abundance of *Klebsiella aerogenes* (beta = 0.041, 95% CI = 1.7 – 2.2, p-value = 0.004) and *Prevotella buccalis* (beta = 0.045, 95% CI = 1.7 – 2.2, p-value = 0.02) (**Figure 4**).

**Figure 4.**
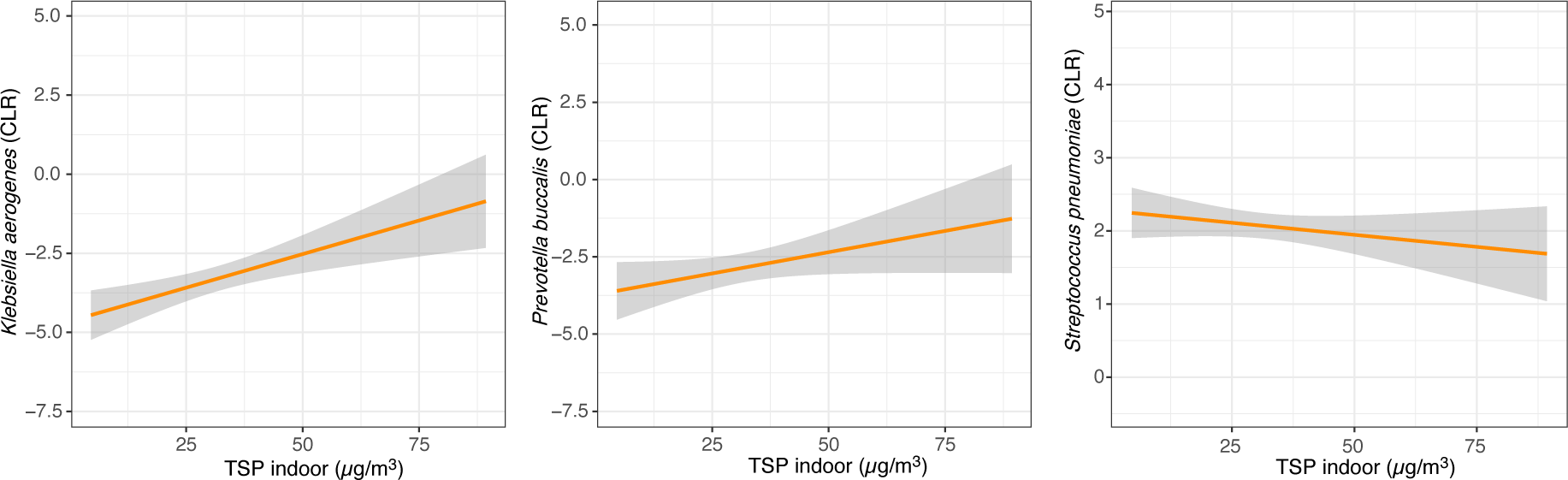
Plots showing significant associations between TSP indoor exposure (µg/m^3^) and bacterial relative abundance (CLR transformed) in the anterior nares of healthy individuals. The model was adjusted for age, gender, days spent in the office, and habitual use of a face mask. The sample observations were grouped by *“subject id”*.

For the NP, indoor TSP exposure was negatively associated with three species of *Corynebacterium* (*C. accolens*: beta = −0.036, 95% CI = 0.9 – 1.3, p-value = 0.03; *C. macginleyi*: beta = −0.033, 95% CI = 0.7 – 1.1, p-value = 0.05; and *C. segmentosum*: beta = −0.033, 95% CI = 0.8 – 1.2, p-value = 0.05), but positively associated with *S. aureus* (beta = 0.012, 95% CI = 0.4 – 0.5, p-value = 0.03) (**Figure 5**).

**Figure 5.**
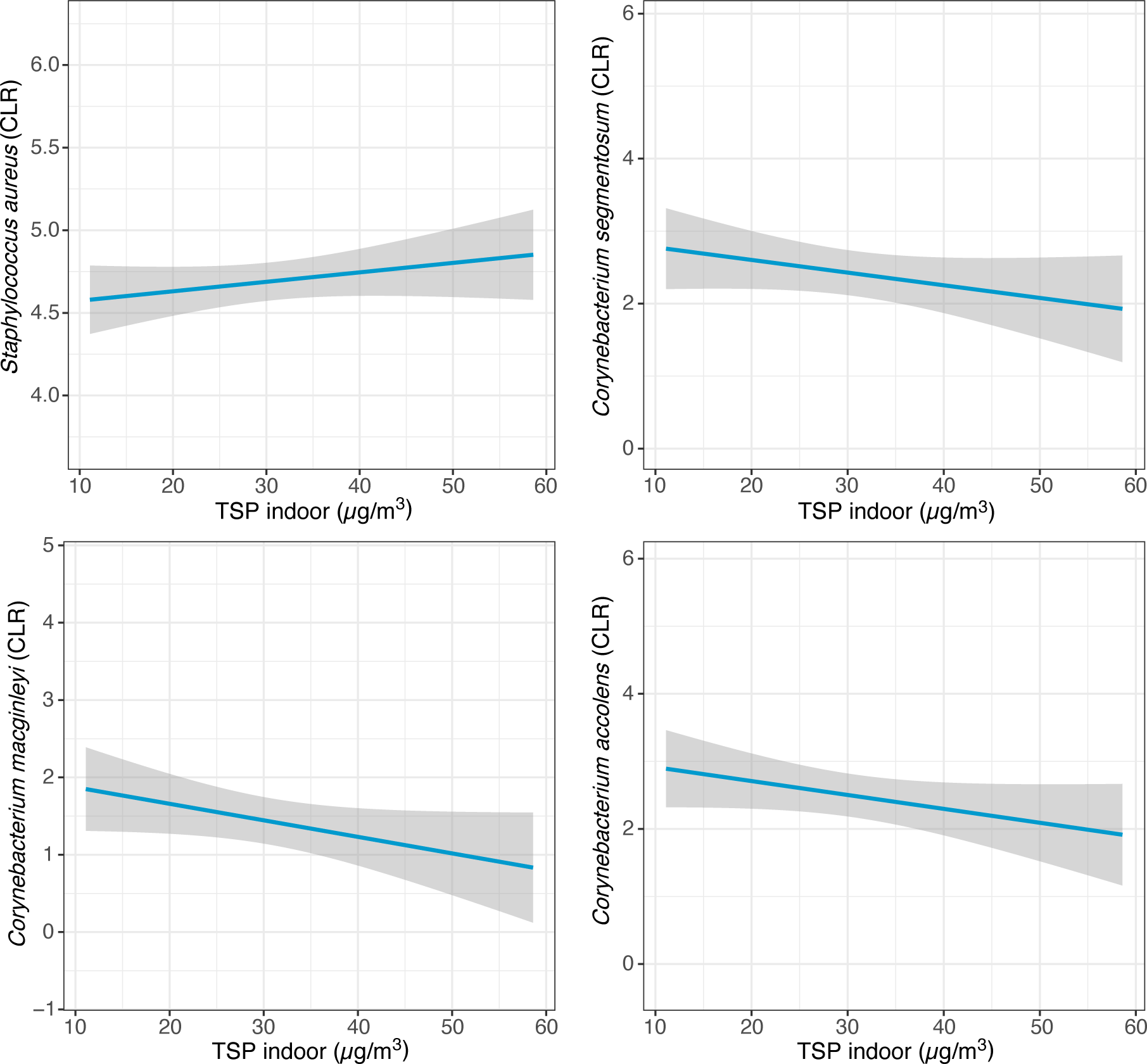
Plots showing significant associations between TSP indoor exposure (µg/m^3^) and bacterial relative abundance (CLR transformed) in the nasopharynx of healthy individuals. The model was adjusted for age, gender, days spent in the office, and habitual use of a face mask. The sample observations were grouped by *“subject id”*.

We also tested for correlations between the microbiome of the indoor TSP samples and the respiratory samples. When comparing with the AN samples, we observed significant correlations in the relative abundance of *S. aureus* (Pearson, rho = 0.25, p-value = 0.04) and *Rothia mucilaginosa* (Pearson, rho = 0.39, p-value = 0.001). When comparing with the NP samples, we observed a significant correlation in the relative abundance of *C. tuberculostearicum* (Pearson, rho = 0.25, p-value = 0.04). For both respiratory samples (AN and NP), we found significant correlations in the relative abundance of *K. pneumoniae* (AN: rho = 0.30, p-value = 0.02; NP: rho = 0.35, p-value = 0.004) and *Cutibacterium granulosum* (AN: rho = 0.31, p-value = 0.01; NP: rho = 0.31, p-value = 0.01) (**Figure 6)**.

**Figure 6.**
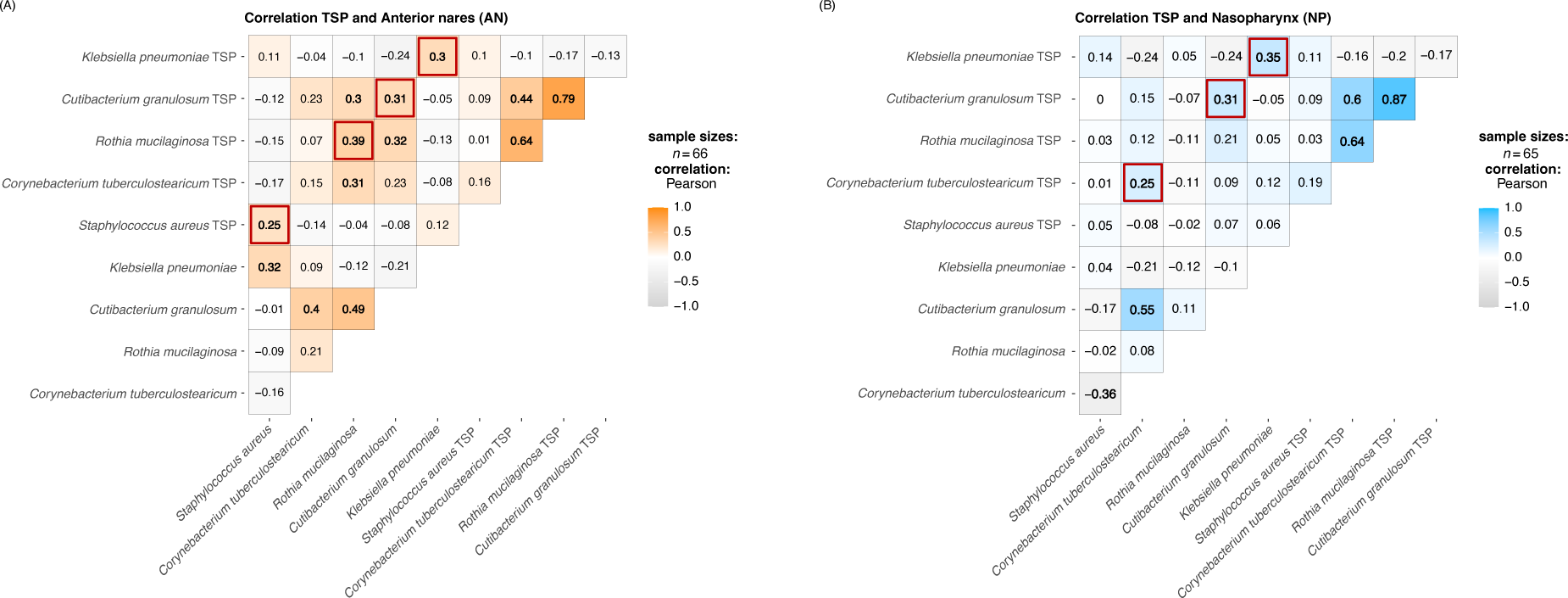
Correlations (Pearson correlation test) in relative abundance between indoor TSP and respiratory samples: anterior nares (AN) and nasopharynx (NP). The red squares highlight the same taxon found in both sample types for which relative abundance is significantly correlated (p-value < 0.05).

We did not observe a significant correlation between antimicrobial resistance gene classes (**Figure 3**) and the alpha diversity of both AN and NP. However, we found a positive correlation between Mercury resistance in the indoor TSP microbiome and Mercury resistance in the NP microbiome (Pearson, rho = 0.44, p-value = 0.0002). Taken together, this suggests a correlation between the indoor TSP microbiome and the upper respiratory microbiome.

## 4. Discussion

Particulate matter has been extensively studied because of its known detrimental impact on human health. However, few studies have analyzed the microbiome associated with these particles, mainly focusing on outdoor exposure of PM_10_ or PM_2.5_ in a limited number of cities (Bertolini et al., 2013; Cao et al., 2014; Gou et al., 2016; Qin et al., 2020). In the present study, we analyzed the microbial composition of indoor and outdoor Total Suspended Particles (TSP) samples collected in Milan and Como, Italy. We investigated the possible association between particle concentration and microbial composition of the human upper respiratory microbiome. Specifically, we monitored 13 offices located in 4 buildings for 3 weeks between May and July 2022. At the end of each week, we collected TSP samples inside the offices and outside the office buildings through a filter-based technique and respiratory samples (anterior nares and nasopharynx swabs) of 26 subjects working in these offices.

Across the three collection periods the indoor TSP was consistently lower compared to the outdoor levels, resulting in indoor/outdoor (I/O) ratios generally <1. The lowest I/O ratios (on average ∼ 0.4) were observed in the third monitoring group for the offices in Como that are located in the basement, which were characterized by very low indoor TSP concentrations, probably because of general building characteristics and construction features. Indeed, while the offices in Milan are located in old buildings with natural ventilation and a heating and/or a cooling system that can result in higher infiltration from outdoors, the buildings in Como are of a recent construction and all the offices had mechanical ventilation and heating and cooling systems. Moreover, the offices in the basement level do not have windows that can be opened, and they are in a clean area, with lower foot traffic.

Besides the difference in concentrations, indoor and outdoor TSP samples differed also in microbiome diversity and composition. The bacteria identified in the TSP samples included mostly soil, plants, and human skin bacteria. Several species of *Paracoccus* (i.e., *P. yeei*) were enriched in the indoor samples. This taxon is likely a commensal that can be associated with human infections, especially in immunocompromised individuals (Daneshvar et al., 2003; Lasek et al., 2018; Mohd-Afzal et al., 2021). In the indoor samples, we found common members of the upper respiratory tract that sometimes can also become opportunistic pathogens, such as *S. aureus* and *K. pneumoniae*. Conversely, the outdoor samples were enriched in some plant pathogens, a consequence of the location of the buildings monitored that are surrounded by urban parks. Among the two types of TSP samples, we also found shared taxa, such as *Micrococcus luteus*, that were nearly ubiquitous across the samples. This bacteria is found in human skin, soil, dust and air (Ahle et al., 2023; Cao et al., 2014; Madsen et al., 2023.), and is sometimes associated with bloodstream infection in hospitalized patients (Zhu et al., 2021). Other bacteria already found in airborne samples and abundant in our samples were *Kocuria rosea, Kocuria rhizophila,* and *Moraxella osloensis* which are common environmental contaminants (Cao et al., 2014; Kooken et al., 2012; Madsen et al., 2023). In addition, both indoor and outdoor particles carried antimicrobial resistance genes. Among the antimicrobial resistance class identified in our data, tetracycline was one of the most common. Tetracyclines are used to treat a variety of infections, and previous studies on pollutants have described their presence in the air raising concerns about the potential transmission capacity of air (Chang et al., 2023; Gao et al., 2023). Overall, we noticed differences in both concentrations of pollutants and microbial composition between the different types of TSP samples. These differences suggested that the outdoor TSP variations only partially influenced indoor TSP. As is well documented, the PM concentration levels in an indoor environment are influenced by multiple variables (Li et al., 2017), including a number of potential indoor sources and factors (e.g., type of ventilation, air exchange rates, type and frequency of indoor activities, number of occupants), and our data suggested that the indoor TSP microbiome is not entirely dependent on the outdoor environment.

We compared the human respiratory microbiome of two upper airway sites, anterior nares and nasopharynx, in healthy subjects. A previous study compared these two types of microbiota using 16S rRNA sequencing data (De Boeck et al., 2017). In our study, we observed significant difference in microbial diversity between the nasopharynx samples and the anterior nares samples. Specifically, the nasopharynx microbiome showed higher diversity than the anterior nares samples. In line with the previous study (De Boeck et al., 2017), we found that the dominant species in the anterior nares were skin bacteria such as *Cutibacterium acnes, S. epidermidis,* and several *Corynebacterium*. Considering that the anterior nares are the most external part of the upper respiratory tract, it is not surprising that some taxa are shared with the skin microbiome (Smythe and Wilkinson, 2023). In our subjects the dominant species in the nasopharynx was *S. aureus*, however other “community types” in the nasopharynx were identified by De Boeck et al. These differences can be a consequence of the human population sampled in the study. Overall, besides these differences, the anterior nares and nasopharynx microbiota appeared to be similar and mostly dominated by gram-positive bacteria.

We analyzed the correlation of the respiratory microbiome and the indoor TSP concentrations. We found that TSP exposure affected the relative abundance of several gram-positive bacteria that are usually commensal in the upper airways, such as the *Corynebacterium sp*. Previous studies have described associations between air pollution and respiratory microbiota, especially in people with respiratory conditions (Chen et al., 2020; Mariani et al., 2021; Vieceli et al., 2023). In our analysis, we also found a positive association between indoor TSP exposure and the relative abundance of *S. aureus*. This is in line with previous cell culture studies that reported that PM and cigarette smoke increased the presence of *Staphylococcus aureus* in samples from the upper airways to potentially pathogenic levels (Lacoma et al., 2019, 2019; Purves et al., 2022).

Finally, the relative abundance of skin and respiratory bacteria identified in the indoor TSP had a positive correlation with the relative abundance of these bacteria in the respiratory samples, suggesting a potential interaction between these air pollutants and our respiratory microbiome. Specifically, we observed significant correlations in both respiratory sites (AN and NP) with the relative abundance of *K. pneumoniae* and *Cutibacterium granulosum*. While *C. granulosum* is generally found in skin and is not associated with infections, *K. pneumoniae* has been widely reported in respiratory infections (Choby et al., 2020; Martin and Bachman, 2018; Wang et al., 2020). We also found a positive correlation with *S. aureus* relative abundance in the anterior nares. Taken together, these findings suggest the TSP microbiome correlates with the upper respiratory microbiome of individuals present in an enclosed environment.

## 5. Conclusion

To the best of our knowledge, this is the first study to characterize the microbiome of indoor TSP, comparing it with the microbiome of outdoor TSP, combined with an analysis of correlations with the human respiratory microbiome. Our results suggest that the indoor TSP has its own microbiome that differs from the outdoors in terms of diversity and composition, and that its composition correlates with TSP concentrations. We also found an association between bacteria identified in the indoor TSP and the upper respiratory tract, indicating that exhaled breath is likely contributing to seeding the indoor TSP. Further analysis in larger populations is needed to validate these preliminary findings and investigate the relationship between these pollutants and the human respiratory tract.

## CRediT authorship contribution statement

**Giulia Solazzo**: writing – original draft, conceptualization, and formal analysis; **Sabrina Rovelli:** resources and data curation; **Matthew Chung**: data curation and software; **Michael Frimpong**: data curation and formal analysis; **Simona Iodice:** data curation and writing – review & editing; **Valentina Bollati** and **Luca Ferrari**: conceptualization, and writing – review & editing; **Elodie Ghedin**: resources, supervision, project administration, and writing – review & editing

## Acknowledgments

This work was funded in part by the Division of Intramural Research (DIR) of the National Institute of Allergy and Infectious Diseases, National Institutes of Health. This work utilized the computational resources of the Office of Cyber Infrastructure and Computational Biology (OCICB) High Performance Computing (HPC) cluster at the National Institute of Allergy and Infectious Diseases (NIAID), Bethesda, MD. Sequencing and initial data processing were conducted at the Frederick National Laboratory for Cancer Research (FNLCR) at the CCR Sequencing Facility, NCI, NIH, Frederick, MD.

## Data availability

The sequencing data from this study are available in the Sequence Read Archive (SRA) under the following accession number: PRJNA1129090.

## Ethics approval

This study was approved by the Ethics Committee of the University of Milan (approval number 24/22), in agreement with the principles of Helsinki Declaration.

