## Supplemental Materials for "The microbiome of Total Suspended Particles (TSP) and its influence on the respiratory microbiome of healthy office workers"

**Supplemental Table 1.** Indoor/Outdoor (I/O) ratios, calculated by dividing each TSP indoor concentration with the average of the corresponding outdoor concentrations. Ratios > 1 are highlighted in red.

| I/O ratios (μg/m$^3$) | | | | | | | | |
|---|---|---|---|---|---|---|---|---|
| **Office** | **Outdoor area*** | **T1** | **T2** | **T3** | **Average of the three weeks** | **St. Dev.** | **Min.** | **Max.** |
| Office 1 | Out1 | 0.9 | 0.7 | 0.8 | 0.8 | 0.1 | 0.7 | 0.9 |
| Office 2 | Out1 | 0.8 | 0.8 | 0.8 | 0.8 | 0.03 | 0.8 | 0.8 |
| Office 3 | Out1 | 1.1 | 1.2 | 1.0 | 1.1 | 0.1 | 1.0 | 1.2 |
| Office 4 | Out2 | 0.8 | 0.6 | 0.6 | 0.7 | 0.1 | 0.6 | 0.8 |
| Office 5 | Out2 | 0.9 | 0.6 | 0.5 | 0.6 | 0.2 | 0.5 | 0.9 |
| Office 6 | Out2 | 1.1 | 0.8 | 0.8 | 0.9 | 0.1 | 0.8 | 1.1 |
| Office 7 | Out2 | 0.9 | 0.6 | 0.5 | 0.7 | 0.2 | 0.5 | 0.9 |
| Office 8 | Out2 | 0.7 | 0.6 | 0.5 | 0.6 | 0.1 | 0.5 | 0.7 |
| Office 9 | Out3 | 0.6 | 0.6 | 0.5 | 0.6 | 0.04 | 0.5 | 0.6 |
| Office 10 | Out3 | 0.7 | 0.6 | 0.5 | 0.6 | 0.1 | 0.5 | 0.7 |
| Office 11 | Out3 | 0.4 | 0.3 | 0.4 | 0.4 | 0.1 | 0.3 | 0.4 |
| Office 12 | Out3 | 0.3 | 0.4 | 0.4 | 0.4 | 0.04 | 0.3 | 0.4 |
| Office 13 | Out3 | 0.4 | 0.4 | 0.4 | 0.4 | 0.01 | 0.4 | 0.4 |

*Out1 indicates the outdoor area of the first monitoring group (MG-1), Out2 indicates the outdoor area of the second monitoring group (MG-2), and Out3 indicates the outdoor area of the third monitoring group (MG-3)

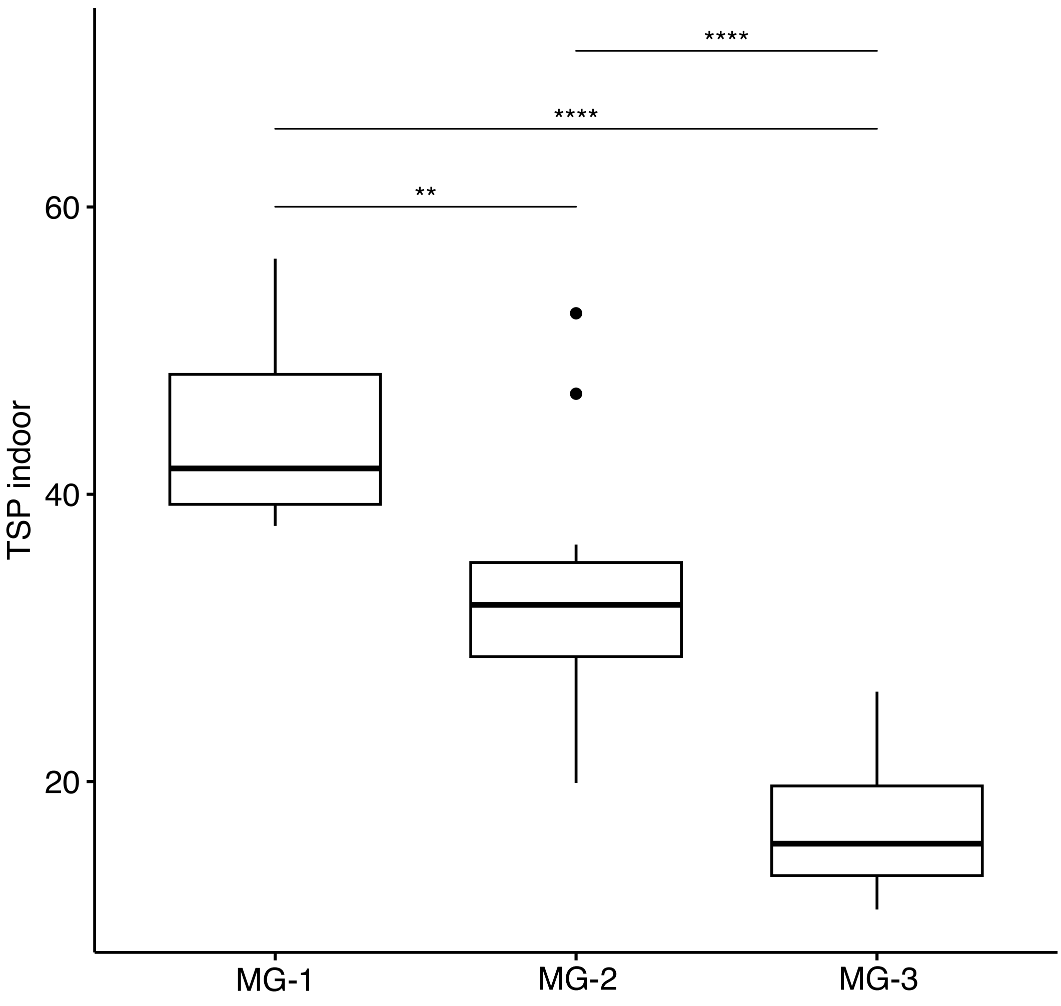


**Supplemental Figure 1.** Indoor TSP concentrations (µg/$m^3$) measured in each monitoring groups (MG-1, MG-2, and MG-3) over the 3 weeks. The stars indicate the significance from the Wilcoxon test (p-value: * < 0.05; ** < 0.01; *** < 0.001; **** < 0.0001).

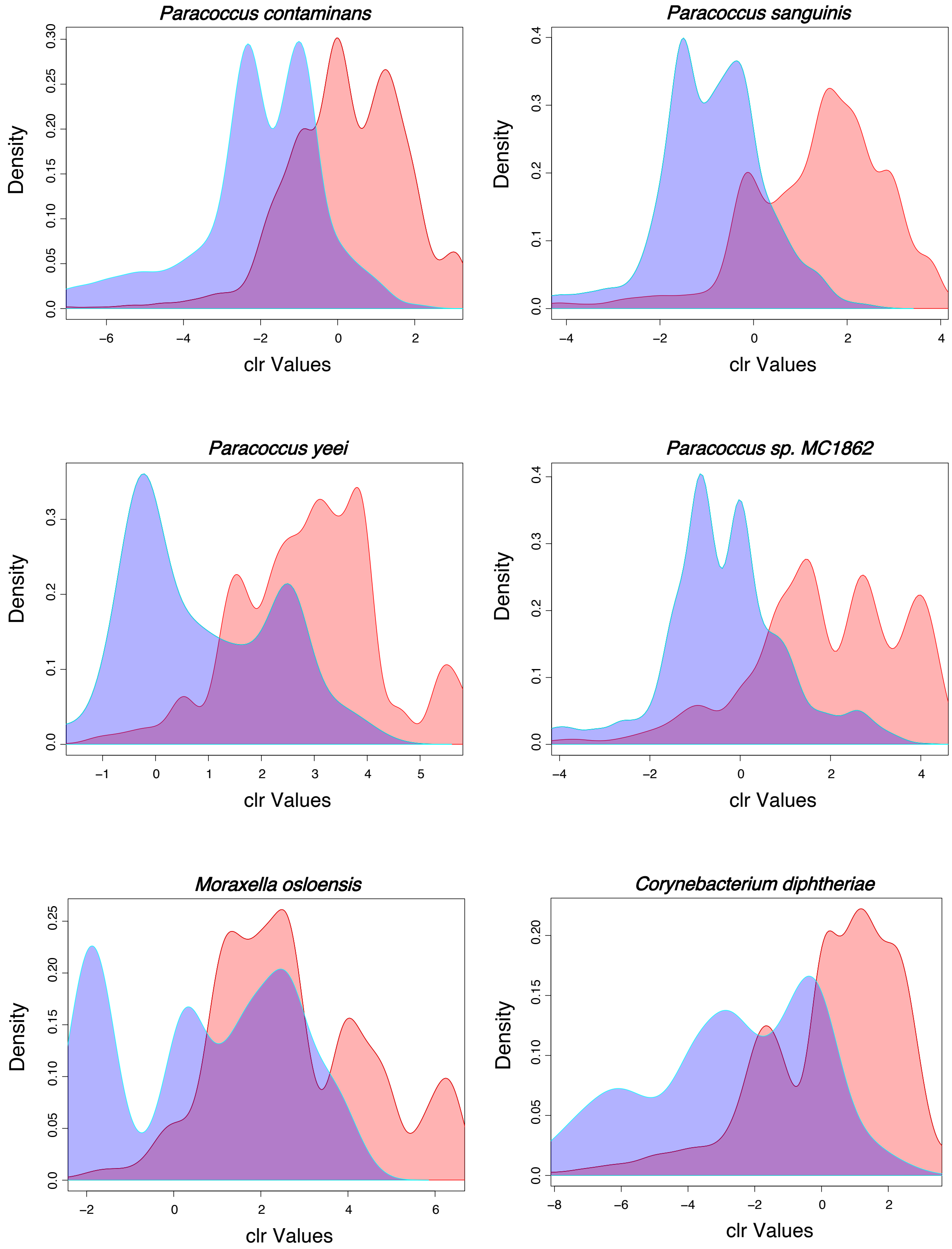


**Supplemental Figure 2.** Centered-log ratio (CLR) transformed abundance of bacteria that are differentially abundant in the indoor and outdoor TSP samples. The red curve represents the indoor TSP samples, while the blue curve represents the outdoor TSP samples. These bacteria were significantly differently abundant (FDR < 0.05) in both ALDEx2 and ANCOM-BC analyses.

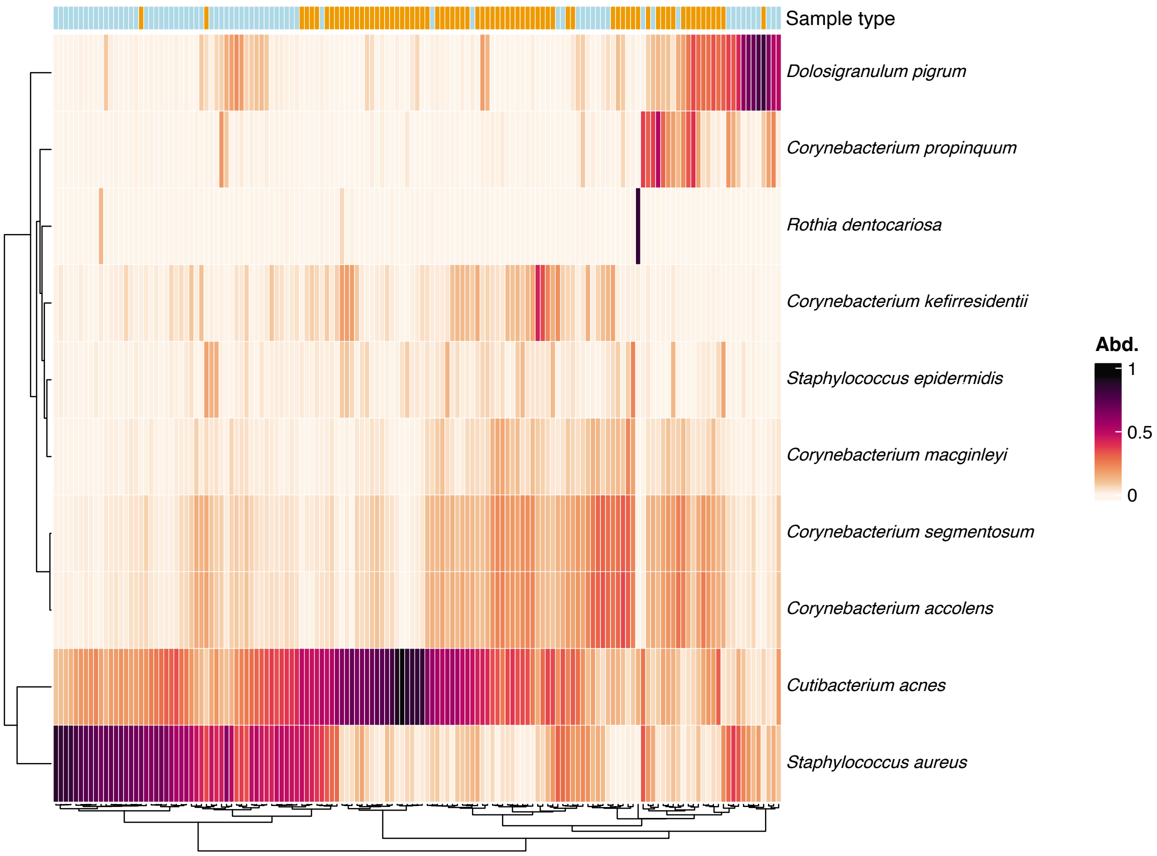


**Supplemental Figure 3.** Heatmap of the relative abundance of the top 10 bacteria identified across all human respiratory samples (sample type), anterior nares samples (orange) and nasopharynx samples (light blue).

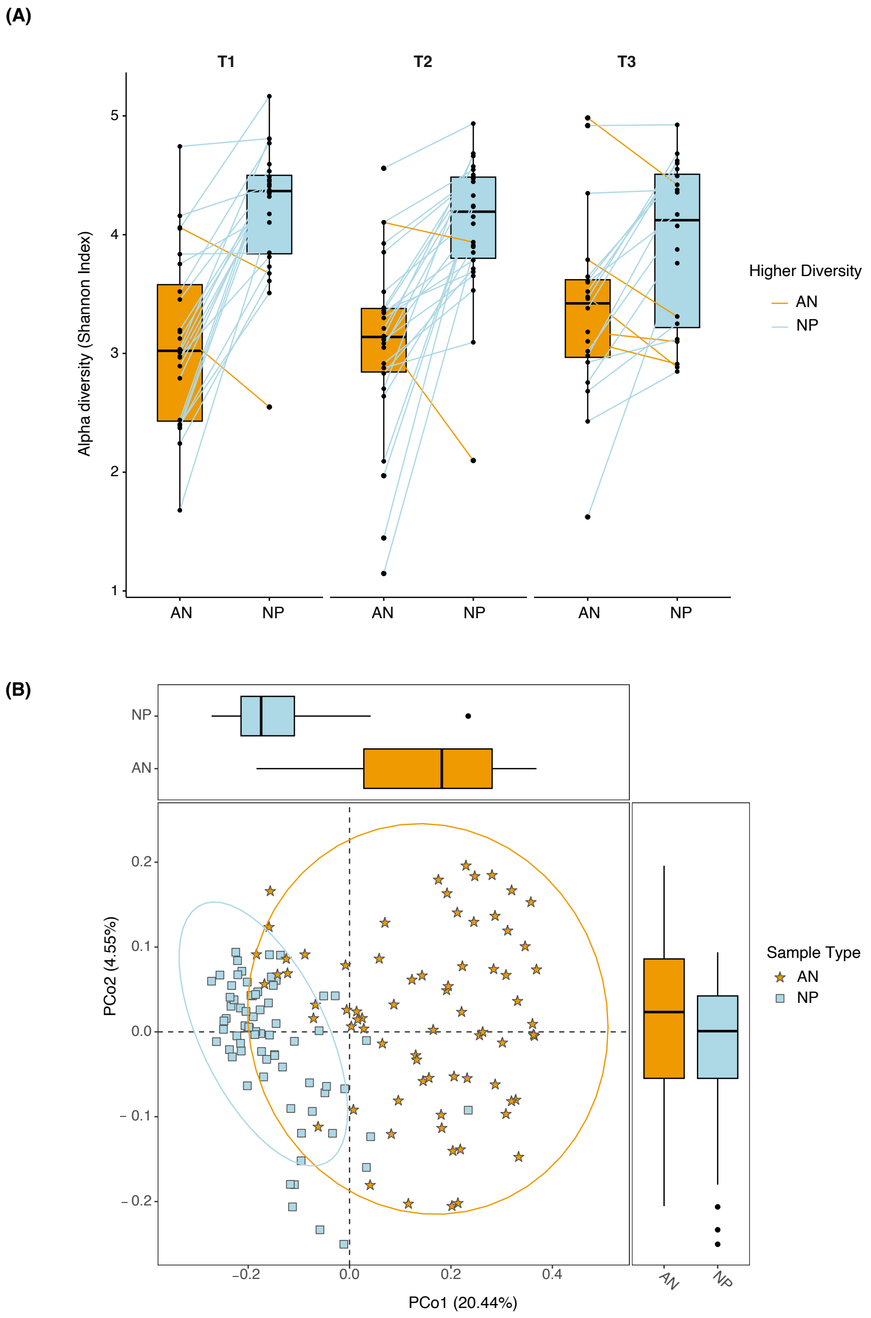


**Supplemental Figure 4.** Microbiome diversity in anterior nares (AN) and nasopharynx (NP) of healthy subjects. (**A**) Alpha diversity comparisons between AN and NP across the weeks: T1 corresponds to the first week of each monitoring group (MG-1, MG-2, MG-3), T2 indicates the second week, and T3 indicates the third week. The line connects the alpha diversity values of the same subjects between the two sampling sites (AN and NP), when the alpha diversity is higher in AN, the line is orange, when the alpha diversity is higher in NP, the line is blue. (**B)** Beta diversity (Bray-Curtis matrix), the orange stars indicates the AN samples and the blue squares the NP samples.

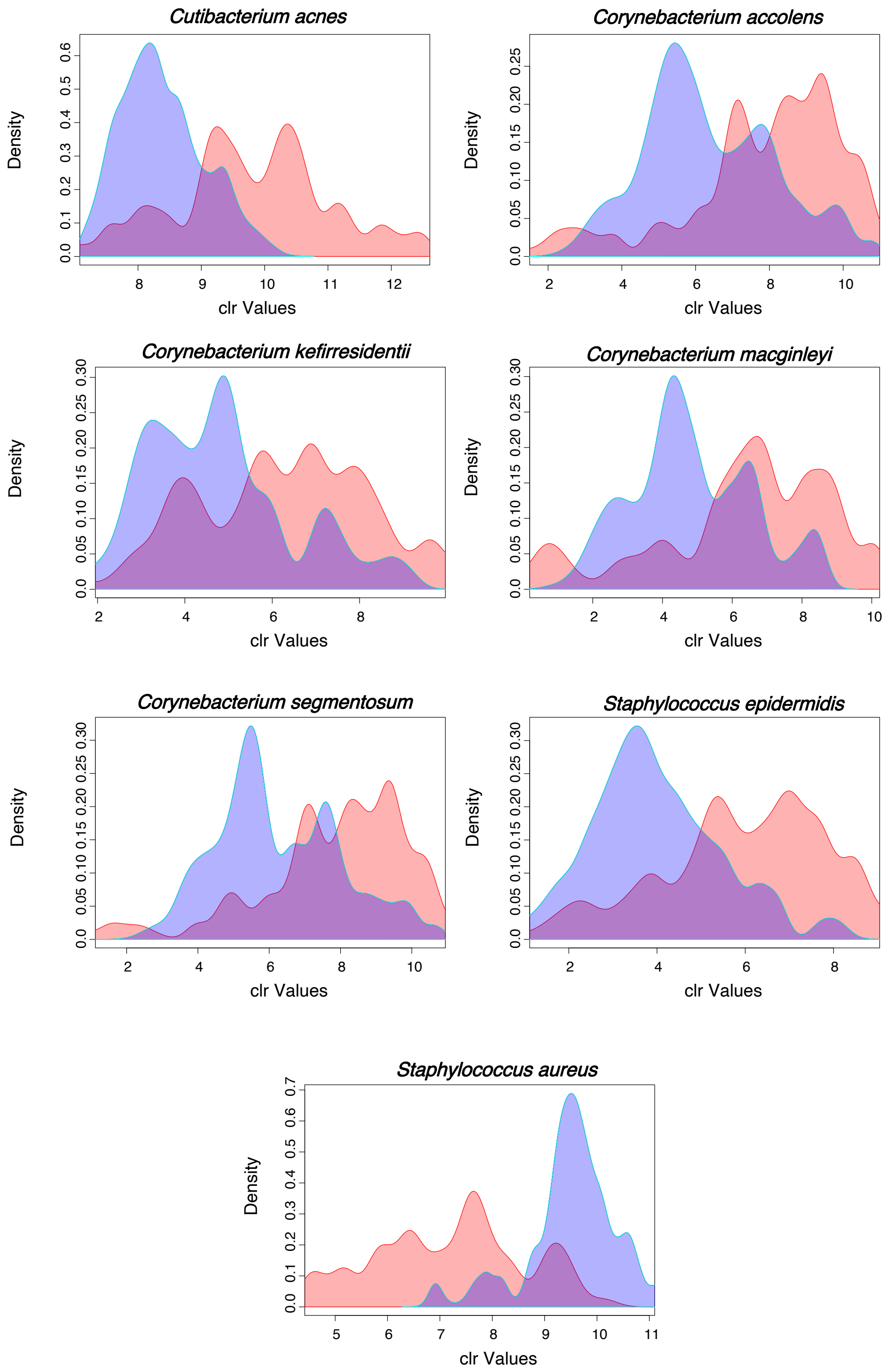


**Supplemental Figure 5.** Centered-log ratio (CLR) transformed abundance of bacteria that are differentially abundant in the Anterior nares (AN) and Nasopharynx (NP) samples. The red curve represents the AN samples, while the blue curve represents the NP samples. These bacteria were significantly differently abundant (FDR < 0.05) in both ALDEx2 and ANCOM-BC analyses.
